# Enrichment of convergent metabolic functions in microbial communities through imposed and emergent environmental niches

**DOI:** 10.64898/2026.02.11.705344

**Authors:** Alberto Scarampi, Sarah J. N. Duxbury, Mary Coates, A. Murat Eren, Orkun S. Soyer

**Affiliations:** School of Life Sciences, University of Warwick, Coventry, UK; Helmholtz Institute for Functional Marine Biodiversity, Oldenburg, Germany; Alfred Wegener Institute, Bremerhaven, Germany; University of Oldenburg, Germany; Max Planck Institute for Marine Microbiology, Bremen, Germany

## Abstract

Microbial community composition is determined by numerous factors, including environmental filtering, biotic interactions, and stochasticity. Disentangling the relative contribution of these distinct processes is a fundamental problem for microbial ecology. Towards addressing it, here we used controlled experiments on replicate freshwater communities that vary individual factors, and used contig-based metagenomics methods to capture functional composition independently from uneven genome recovery across taxa. After repeated sub-culturing in minimal media lacking organic carbon but containing nitrate and vitamins, communities with distinct initial compositions consistently converged toward similar taxonomic structures and metabolic functions. These included oxygenic photosynthesis and nitrate metabolism, consistent with the imposed growth regime, and also showed reproducible enrichment of anoxygenic photosynthesis, vitamin biosynthesis and degradation pathways. These patterns indicate strong environmental filtering during assembly while also revealing a consistent role for emergent environments arising from microbial activity and metabolic interactions. To more directly test specific environmental drivers, we initiated replicate cultures from a single community and propagated them in the original medium and in variants lacking vitamins or both vitamins and nitrates. Only nitrate removal produced a distinct and statistically significant shift in both composition and function, with nitrate-free communities enriched for cyanobacterial nitrogen fixation and supporting metabolic functions in heterotrophs. Together, these results support a hierarchy of environmental filters determining community outcomes and provide a quantitative framework for predicting and steering community function through rational environmental design.

## Introduction

Microbial communities underlie Earth’s biogeochemical cycles [1] and are integral to host biology [2], yet which processes control their taxonomic and functional composition remains an open question in microbial ecology [3, 4, 5]. Both deterministic forces (environmental selection, niche differentiation) and stochastic processes (dispersal, drift, historical contingency) can result in stable composition and possible alternative compositional states in communities [3, 4, 2]. Resolving when and how these processes dominate community assembly is critical to understanding the design principles of complex microbial communities.

Comparative metagenomics studies across distinct environments revealed environment-specific functional composition [6, 7], where the metabolic modules encoded in the community correlate with environmental factors [8]. Critically, functional composition seems to correlate more strongly with environmental factors than taxonomic composition [9], and functional repertoires often seem to persist despite taxonomic differences [10, 11, 12, 13]. Persistent and specific functional composition is also observed in host- and particle-associated communities [14, 11, 15, 16], even though stochastic colonization dynamics are expected to influence initial composition in these cases. While these observations implicate environmental factors as important determinants of functional composition, the lack of causal inferences and mechanistic insights into how specific environmental factors influence function limits the translation of such observations into predictive frameworks for how communities respond to environmental change and into rational strategies for engineering stable consortia.

Laboratory enrichments of natural microbial communities under defined conditions provide a more direct route to test causal links between environment and community function while removing confounding processes such as predation and immigration [17]. Enrichment studies have shown reductions in species richness and varying degrees of convergence in taxonomic composition [18, 19]. Many of these studies found that communities maintained relatively high diversity despite well-defined, simple, and stable environmental conditions, suggesting a role for within-community interactions and the emergence of new metabolic niches. For example, in enrichments from soil isolate mixtures, communities grown under abundant glucose showed cross-feeding interactions [18, 19], and the extent of diversity correlated with glucose concentrations or the number of available carbon sources [20, 21, 22]. In enrichments from freshwater communities, and without added carbon source, final communities repeatedly showed specific metabolic functions including (an)oxygenic photosynthesis, polysaccharide degradation, and vitamin biosynthesis [23, 24]. Overall, these studies suggest a role for within community interactions, and indicate that environment can directly influence community composition. However, a detailed, mechanistic understanding of environment-function relation is lacking due to many studies having been limited to taxonomic surveys and not being able to track reproducible selection of specific metabolic functions between source and enrichment communities [18, 20, 21].

Altogether, both environmental genomic studies and laboratory enrichments highlight the crucial role of the environment in shaping community functional repertoire. However, key questions remain unresolved: Does environment drive community composition deterministically, resulting in repeatable community compositions? If so, which metabolic functions are consistently selected? Do certain environmental factors exert stronger selective pressure than others to dictate functional outcomes? Here, we address these questions using freshwater photosynthetic communities, which offer tractable and ecologically relevant model systems for environment-community relation [25, 24, 26, 27, 23]. We perform enrichments from different starting communities in minimal media without carbon source and with nitrate, and from the same starting community under varying environmental conditions. We complement our experimental workflow with comprehensive characterizations of microbial diversity and metabolism that employ genome-resolved as well as assembly-level analyses of shotgun metagenomes to gain high-resolution insights into microbial succession and enrichment of metabolic modules. Our findings show that different starting communities converge to a common final community composition in a repeatable manner, when grown in minimal media without carbon source and with nitrate. Final communities show reproducible enrichment of (an)oxygenic photosynthesis and nitrate metabolism pathways. In addition, assembly from the same starting community is deterministically altered in the absence of nitrate, but not to the same extent in the absence of vitamins, indicating that nitrate availability exerts stronger selective pressure than micronutrient supplementation on functional community composition. Overall, these findings show that controlling specific environmental factors under stable conditions can deterministically alter community function.

## Results

### Laboratory enrichment leads to reduced diversity and enhanced community stratification

To investigate how complex natural communities adapt to defined laboratory conditions, we initiated eight cultures from freshwater samples collected from Draycote Reservoir at different times over a 2-year period (see *Methods*). Each culture was maintained in minimal medium lacking organic carbon and containing nitrate, under 12-hour light-dark cycles with sub-culturing (see Methods and Fig. S1). In most enrichment cultures the presence of spatial structures was observed (Figure 1A). After multiple passages, we performed deep metagenomic sequencing on both the original lake samples (“source”) and the laboratory-enriched communities (“enrichment”) to track community changes.

**Figure 1.**
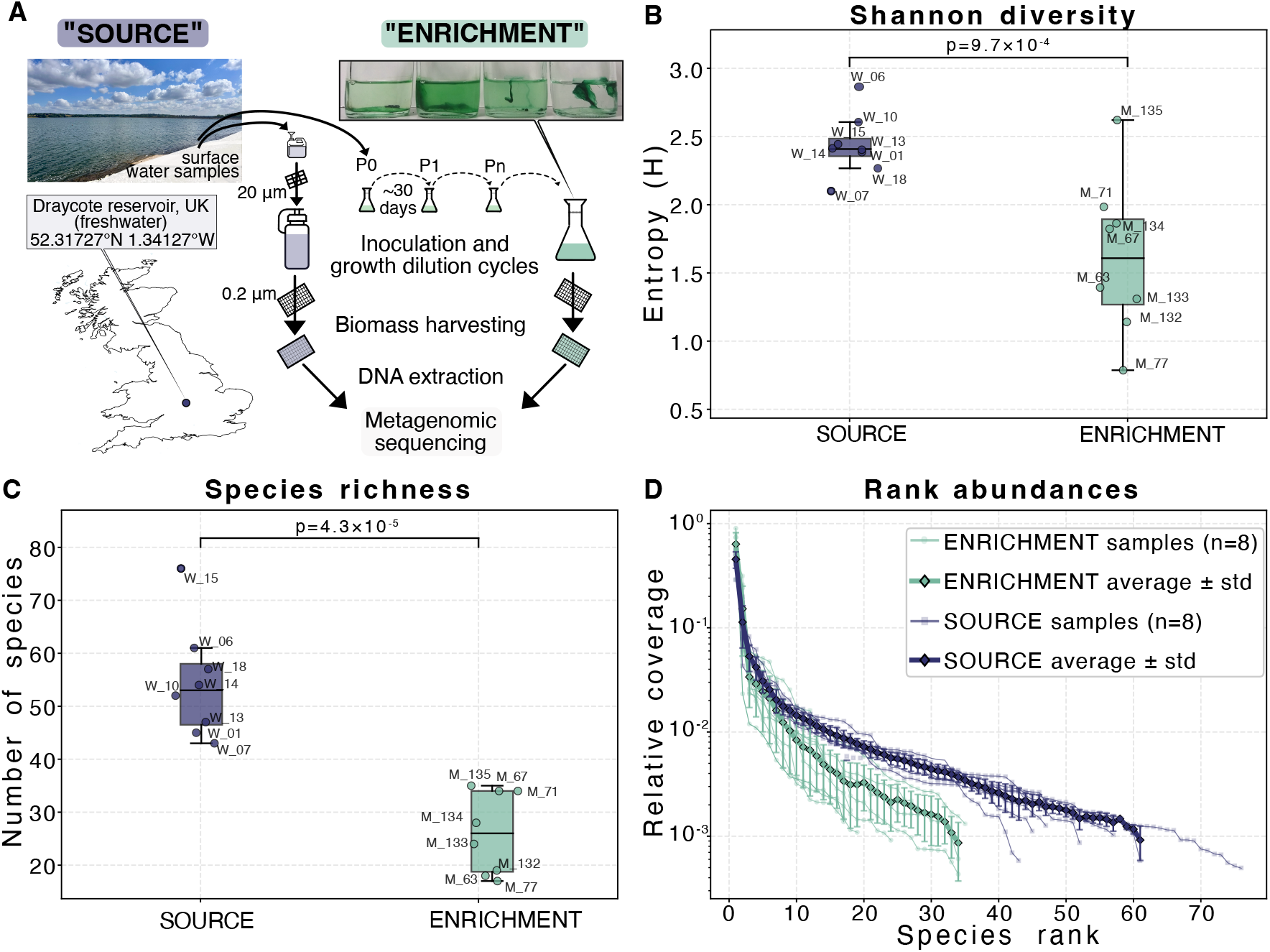
(A) Schematic of sampling and enrichment strategy. Surface water was collected from Draycote Reservoir (52.317°N, 1.341°W, UK), sequentially filtered (200 *µ*m, 20 *µ*m, and 0.2 *µ*m), and used for metagenomic extraction of SOURCE communities and to inoculate BG11+ medium enrichments maintained for >1 year under semi-natural controlled conditions (30^*°*^C; 12/12 h light/dark cycles; 40 *µ*E; 1:50 dilution at each subculturing passage) (see further details in *Methods*). (B) Shannon diversity (Entropy, H) of source water (W) and enrichment (M) samples, showing reduced alpha diversity following long-term enrichment.(C)Population numbers, measured as the number of unique ribosomal protein L19 (RplS) sequences. (D) Rank–abundance distributions illustrating relative coverage of RpL19 sequences across source and enrichment communities (individual samples, and averages ± standard deviation for n=8 replicates each). *** indicates statistical significance at p < 0.001 (pairwise t-test).

Metagenomic sequencing of the replicate source and enrichment samples usually yielded *≈* 10^8^ paired reads per sample, which were analysed using different bioinformatic pipelines (see *Methods*). All samples from a given culture were co-assembled and genomes were recovered using STRONG [28], resulting in 444 high-quality metagenome-assembled genomes (MAGs) across all samples. Genome length distributions were similar across all samples, with most MAGs ranging between 2-6 Mb (Figure S2(B)). Assembly quality metrics demonstrated high completeness (>80%) and low redundancy values (*<*10%) for the majority of MAGs across all samples, indicating successful MAG recovery (Figure S1(C,D)). We used highquality MAGs to recruit short reads from each metagenome to profile the relative abundance of genomes across our samples (see *Methods*) (Figure S3).

In addition to the MAGs we were able to recover, we characterized microbial diversity in our samples using ribosomal protein sequences as phylogenetic markers captured in assembled contigs with EcoPhylo[29] (see *Methods*). We found consistently higher number of taxa in source samples compared to all enrichment samples (Figure 1C). Similarly, Shannon diversity indices were elevated in the source sample compared to enrichment samples (Figure 1B). These patterns were significant across all enrichment samples, indicating reproducible convergence towards simpler communities during laboratory adaptation. Rank abundance curves (Figure 1D) showed a structural difference between communities. The source samples exhibited a gradual decline in relative abundance with rank, characteristic of an even species distribution. In contrast, enrichment samples from the final sub-culture showed an increased steepness in rank abundance curve slope, indicating the emergence of few dominant taxa during laboratory cultivation (Figure 1D). The top three species in enrichment samples accounted for 60-80% of total abundance, compared to only 20-30% in source samples. This shift from evenness to dominance suggests that laboratory conditions impose strong selective pressures favouring specific species and metabolic strategies, which we analysed next.

### Repeated enrichments result in reproducible taxonomic convergence, down to species level

Having established that laboratory enrichment consistently reduces diversity, we next sought to understand whether this simplification resulted in predictable community compositions. To quantify community composition across samples, we initially analyzed taxonomic distributions of metagenome-assembled genomes (MAGs). However, we found this approach to be limited in resolving compositional differences due to differential binning success into MAGs across taxa, resulting in under-representation of low abundance taxa (Figure S3). Quantifying community composition using RpL19, a universal single-copy core gene, via the EcoPhylo workflow helped us overcome this limitation by capturing populations that were present in the metagenome but did not assemble into high-quality MAGs.

Our analysis of RpL19 genes revealed reproducible patterns of taxonomic convergence, where all eight enrichment cultures, despite originating from temporally distinct source samples, converged toward similar taxonomic compositions (Figure 2A). Source communities maintained high coverage across diverse phylogenetic branches, particularly within Actinomycetota and various Pseudomonadota lineages. In contrast, enrichment samples showed concentrated coverage in specific clades: primarily Cyanobacteriota (particularly lineages related to *Limnocylindrus*, WH5701, and JAAUUE01), select Gammaproteobacteria (including *Fonsibacter* and ATX-6F1), and reduced representation across most other phyla. To visualise this taxonomic convergence better, we summarised species convergence at the phylum level (Figure 2B). Source samples exhibited even distribution across 10-12 phyla, with Actinomycetota (30-40%), Pseudomonadota (20-30%), and Bacteroidota (10-15%) as major constituents. All enrichment samples, however, converged to a characteristic profile dominated by Cyanobacteriota (40-60% relative coverage) and specific Pseudomonadota lineages (20-30%), while Actinomycetota, despite their dominance in source samples, were present but significantly reduced (*<*5%) across all enrichments). Principal component, using abundances, further confirmed the statistical significance of this convergence. At the phylum level, PC1 explained 97.3% of variance and completely separated enrichment from source samples (p < 0.001, PERMANOVA), with all eight enrichment replicates forming a tight cluster (Figure 2C).

**Figure 2.**
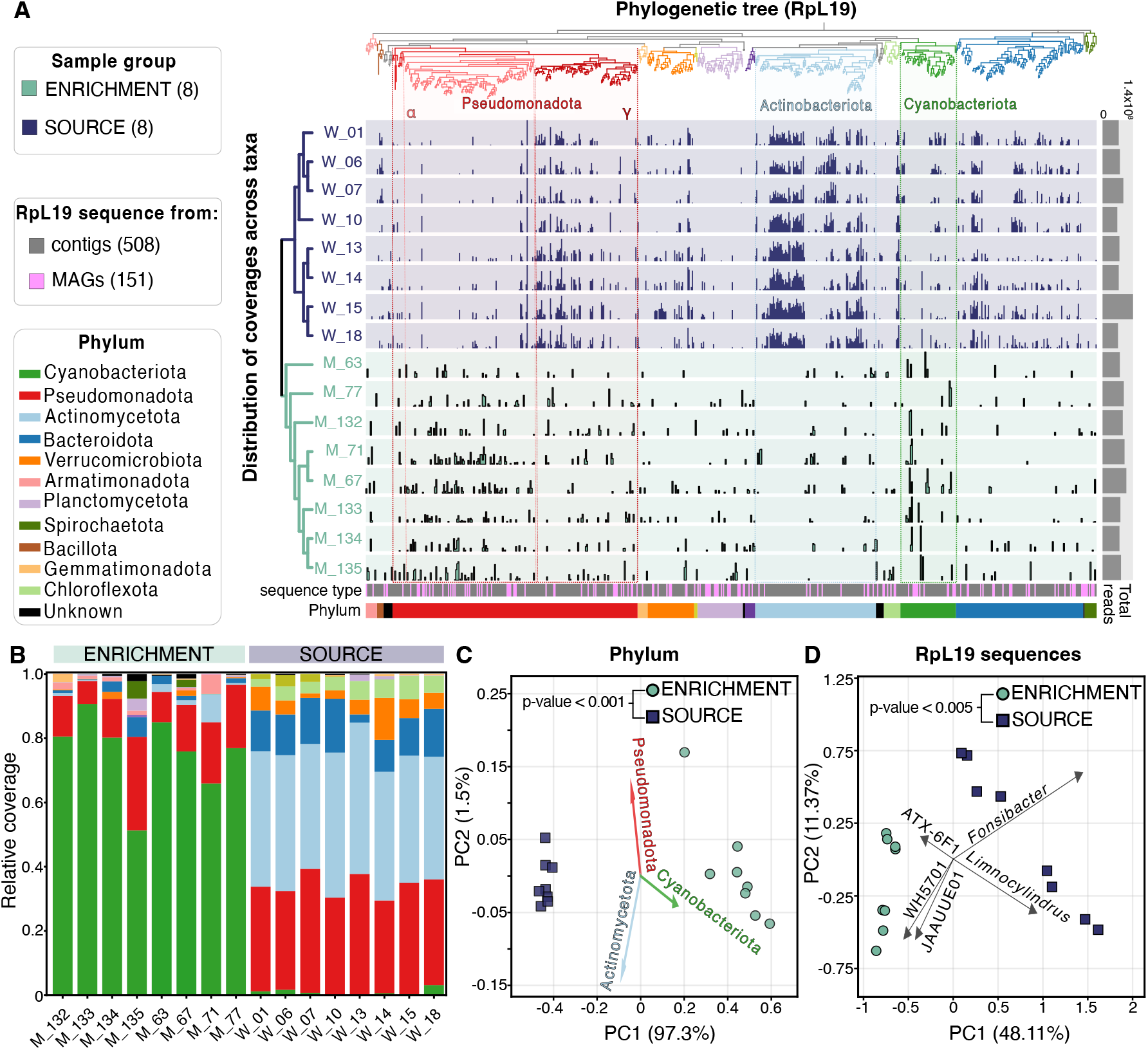
A) Abundance of individual RpL19 sequences across source (labelled W and back-colored blue) and enrichment (labelled M and back-colored light green) communities, summarized using the ecophylo workflow. The dendograms to the left and top of the figure show the community sample clustering and phylogenetic relations, respectively. Total reads in each community sample is shown to the right of the figure, while the bottom of the figure lists the type of the RpL19 sequence (i.e. obtained from a contig or a MAG), and its phylum assignment. The phylogenetic tree is color-coded according to phyla (see legend), and abundant phyla are indicated by name. Some of the individual RpL19 sequences, showing clear differences among source and enrichment communities are highlighted with a thick line. B) Relative abundance of RpL19 sequences, grouped at phylum level and color-coded according to the legend, across all samples. Each set of stacked bars represent an individual enrichment or source community, as indicated on the x-axis. (C, D) Principal component analysis (PCA) of community taxonomic composition at the phylum (C) or individual RpL19 sequence (D) level. Each point represents a source or enrichment community, coloured as in panel A.

To test whether convergence extended to finer taxonomic scales, we repeated this analysis by assigning taxonomy of RpL19 genes at progressively lower taxonomic ranks. While clear separation persisted at class and order levels, the convergence signal progressively weakened at family and genus levels, with variance explained by PC1 dropping from 97.3% at phylum to approximately 40% at genus level. While this observation could indicate that community assembly might be stochastic at the species level, it is also possible that the reduction in the distinction power relates to loss of taxonomic assignment of RpL19 gene as we go down to species levels. When we analyzed clustering patterns using directly the coverage of RpL19 sequences, without taxonomic assignment of those sequences, we recovered strong convergence signals for individual RpL19 sequences (Figure 2D). In this case, PC1 explained 48.11% of variance with clear separation between enrichment and source samples (p < 0.005). This shows that the apparent loss of convergence at lower taxonomic ranks was an artifact of classification challenges: as taxonomic assignment becomes increasingly difficult at finer scales, more sequences fall into “unknown” categories and fail to contribute to the convergence signal. Indeed, we found that *≈* 30% of taxa were unclassified at species level, compared to only < 1% at order level (Fig. S4). The coverage-based analysis that employs *de novo* sequence clusters circumvents this limitation by comparing communities based on actual sequence abundances rather than taxonomic labels.

### Functional analysis reveals metabolic specialization in converged communities, aligned with environmental factors

To test whether taxonomic convergence reflected functional convergence, we analyzed the metabolic capacity encoded in each community using two complementary module-level metrics derived from raw contigs (see Figure 3A and *Methods*). First, we quantified completeness-weighted coverage of KEGG metabolic modules, which captures the abundance of each complete pathway (“abundance-based” metric). Secondly, we estimated the expected copy number of each module per metagenome by calculating stepwise module copy numbers in each metagenome and normalizing them by the number of distinct ribosomal protein marker sequence clusters (count-based metric). This second measure is designed to capture modules, whose count (not coverage) is enriched within the community but they are not necessarily carried out by the most abundant taxa [30].

**Figure 3:**
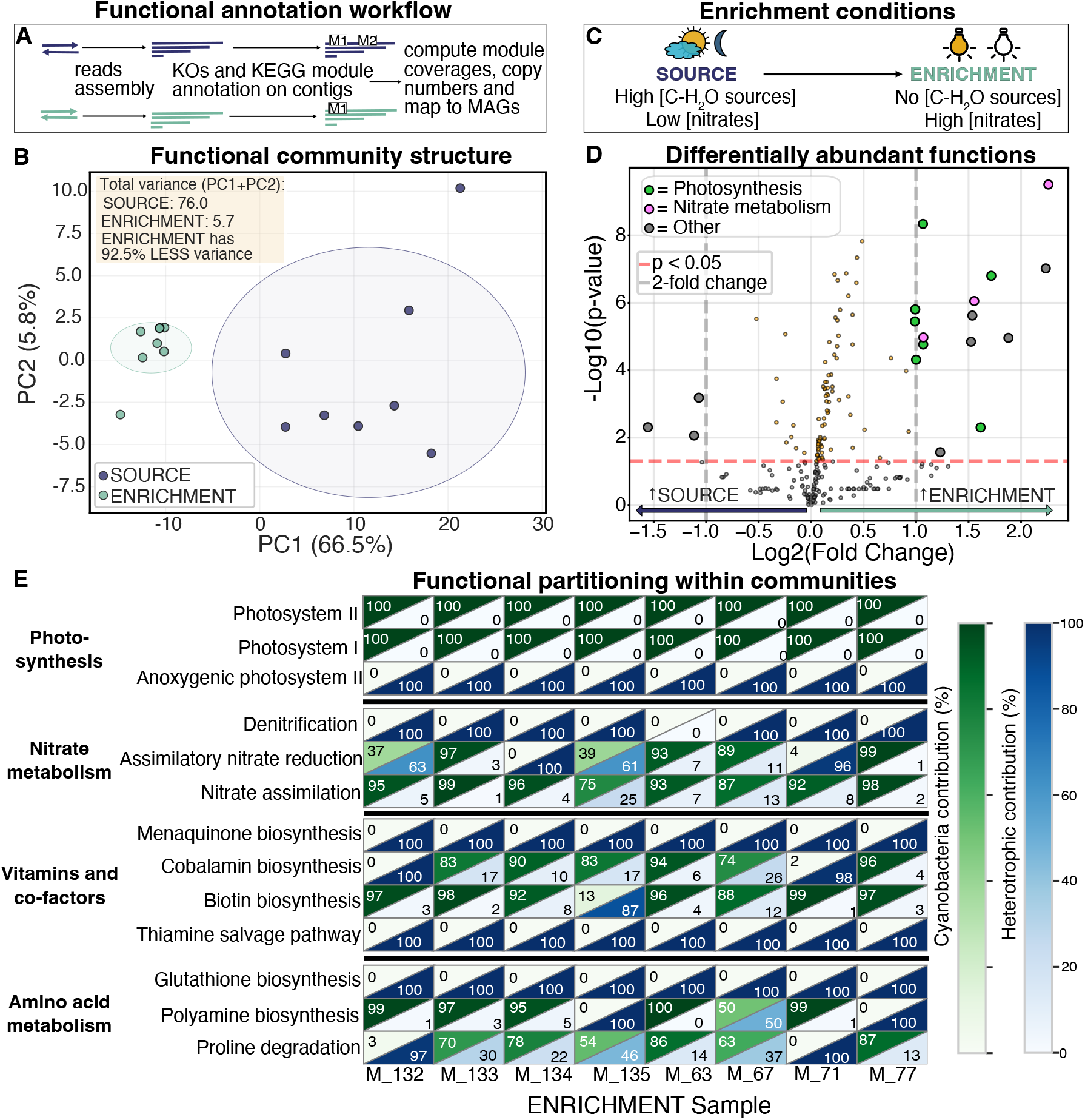
Deterministic functional convergence in enrichment communities. (**A**) Schematic overview of the functional analysis workflow. Metagenomic reads from source (W) and enrichment (M) communities were assembled, ORFs were annotated with KEGG Orthology (KO), and KEGG Modules were reconstructed on raw contigs. For each sample and module we computed abundance-based and (ii) per-genome metrics. See main text and *Methods* for further details. (**B**) Principal component analysis of abundance-based metric for modules across source (blue) and enrichment (green) communities, each shown as a single point. Ellipses indicate 95% confidence regions for each group. Further statistics are shown as inset (**C**) Conceptual summary of environmental conditions during enrichment. Lake-sourced freshwater communities, subject to variable conditions, are inoculated into a carbon-free, nitrate-replete medium under 12h light–dark cycle. (**D**) Differential analysis of metabolic module abundance-based metric between source and enrichment communities. Volcano plot shows log_2_ fold changes in the module abundance-based metric (enrichment/source) versus *−* log_10_,FDR-corrected *p*-values. Each point in the plot shows a module. Modules that are most significantly changed between the enrichment and source communities (FDR < 0.05, red dashed line, and |log_2_FC| > 0.5, gray dashed lines) are shown as bigger dots, with those relating to photosynthesis and nitrogen metabolism pathways colored in green and magenta, respectively. (**E**) Encoding of specific metabolic functions in cyanobacteria vs. all other bacteria (“heterotrophs”) shown as percentage across enrichment communities. Split cells show the relative contribution of cyanobacteriaversus heterotrophs, quantified in green and blue, respectively.

We found that both metrics differed significantly across enrichment and source samples, with specific metabolic pathways showing consistent and significant increase in enrichment communities (Figure 3C,D). PCA on module relative abundances (abundance-based) showed stronger separation of source and enrichment than taxonomy-based analyses, with PC1 explaining 66.5% of variance (PERMANOVA R^2^=0.757, F=43.67, p*<*0.001). Replicate enrichments exhibited a pronounced reduction in variance (92.5% reduction in PC1+PC2 spread), indicating convergence to similar metabolic functional profiles (Figure 3B).

The nature of increased modules, when using the abundance-based metric, matched the imposed environmental conditions: a carbon-free, nitrate-replete, light–dark regime (Figure 3B). Accordingly, the increased modules included: dissimilatory nitrate reduction (M00530), oxygenic photosystems I (M00163) and II (M00161) and associated electron transport components of NAD(P)H:quinone oxidoreductase (M00145) and cytochrome b6f complex (M00162) (Table1). Consistent with autotrophic metabolism, abundance of Calvin cycle pathways was elevated (Fig. S6-S7). In addition, and less clearly deducible from imposed conditions modules involved in vitamin metabolism and antioxidant capacity were prominent: thiamine salvage (M00899), tocophero biosynthesis (M00112), phylloquinone (vitamin K1) biosynthesis (M00932), and anaerobic cobalamin biosynthesis (M00924). Ectoine degradation (M00919) was also high in enrichment communities (Table 1, Fig. S6-S7).

**Table 1:** Top 10 KEGG modules that are increased in enrichment communities. Modules identified using coverage-based and count-based metrics. All of the modules shown display significantly increased metrics in enrichment communities compared to the source communities (p < 0.05).

| Module | Module name | log<sub>2</sub>FC | p<sub>adj</sub> |
| --- | --- | --- | --- |
| *Increase by coverage* |  |  |  |
| M00145 | NAD(P)H:quinone oxidoreductase | 5.67 | 0.002 |
| M00162 | Cytochrome b<sub>6</sub>f complex | 5.51 | 0.026 |
| M00163 | Photosystem I | 5.21 | <0.001 |
| M00530 | Dissimilatory nitrate reduction | 5.02 | <0.001 |
| M00161 | Photosystem II | 4.60 | <0.001 |
| M00899 | Thiamine salvage pathway | 4.52 | 0.001 |
| M00112 | Tocopherol/tocotrienol biosynthesis | 4.10 | 0.017 |
| M00919 | Ectoine degradation | 4.01 | 0.003 |
| M00932 | Phylloquinone biosynthesis | 3.86 | 0.002 |
| M00924 | Cobalamin biosynthesis (anaerobic) | 3.74 | 0.002 |
| *Increase by count* |  |  |  |
| M00091 | Phosphatidylcholine biosynthesis | 0.42 | 0.002 |
| M00854 | Glycogen biosynthesis | 0.35 | 0.002 |
| M00021 | Cysteine biosynthesis | 0.33 | 0.002 |
| M00627 | β-Lactam resistance (Bla system) | 0.33 | 0.002 |
| M00083 | Fatty acid elongation | 0.32 | 0.002 |
| M00118 | Glutathione biosynthesis | 0.32 | 0.002 |
| M00880 | Molybdenum cofactor biosynthesis | 0.26 | 0.004 |
| M00156 | Cytochrome c oxidase (cbb<sub>3</sub>-type) | 0.26 | 0.002 |
| M00970 | Proline degradation | 0.24 | 0.004 |
| M00004 | Pentose phosphate pathway | 0.24 | 0.002 |

The count-based, per-genome metric revealed additional modules that were increased in the enrichment communities, including those involved in: lipid and carbohydrate biosynthesis (phosphatidylcholine biosynthesis, M00091; fatty acid elongation, M00083; glycogen biosynthesis, M00854), amino acid and redox metabolism (cysteine biosynthesis, M00021; glutathione biosynthesis, M00118; proline degradation to glutamate, M00970), central carbon flow (pentose phosphate pathway, M00004), respiratory adaptation (low-pxygen cbb3-type cytochrome c oxidase, M00156), and cofactor assembly (molybdenum cofactor biosynthesis, M00880) Table1. Since the per-genome metric is less biased towards higher abundance members of the community, the increased modules captured by this metric could indicate persistent interactions with or between lower abundance members of the enrichment communities.

Together, increased modules identified through both metrics indicate a deterministic functional shift under our enrichment regime: oxygenic photosynthesis and Calvin cycle for carbon acquisition; nitrate-linked energy and nitrogen transformations; and widespread vitamin/cofactor metabolism and oxidative stress handling. We also examined module variability across enrichment communities. Modules that are significantly increased in enrichment communities, compared to the source communities, showed lower coefficients of variation and were mostly highly abundant (Fig. S8). Only few of the significantly increased modules displayed the opposite trend and were low-abundance (mean relative abundance: 0.047). This shows that increased metabolic functions were consistently increased across all replicate enrichments, corroborating the power of functional profiles to discriminate source and enrichment communities (Figure 3B).

### Phylogenetic distribution of metabolic functions indicate flexibility in community compositions for achieving same functional outcomes

We next analysed the phylogenetic distribution of the modules increased in enrichment communities, i.e. which genomes they are encoded in. While we have performed this analysis at the species and phylum levels (Fig S6,7), we summarize it here with regards to whether a functional module is encoded in cyanobacteria or any other bacteria, broadly referred as “heterotrophic”, as this provides a useful, visual coarse-graining (Figure 3E).

We found, as expected, that the increased oxygenic photosynthetic modules (e.g. Photosystem I, Photosystem II) were invariably encoded exclusively by cyanobacteria across all eight enrichment communities, while denitrification was consistently encoded in heterotrophs (Figure 3E). Almost all other highly increased metabolic functions, however, showed variable patterns across different enrichment communities. For example, biotin biosynthesis was predominantly encoded in cyanobacteria in most communities (e.g., M 63: 99% cyanobacterial), but in one (M 135), it was found in heterotrophs (63% heterotrophic) (Figure 3E). This suggest that a function can be maintained differently in different communities, being encoded in - or contributed by - different taxa. Similar compensatory patterns in module encoding were observed for thiamine salvage, cobalamin biosynthesis, and nitrogen metabolism pathways (Figure 3E). This supports the notion that communities “assemble” to maintain or fulfil certain functions, which can be established by different taxa, or in other words metabolic functions are selected to fill specific environmental niches that can be filled by either cyanobacteria or heterotrophs depending on the specific strain composition of each converged community. In specific taxon-level analyses of module encoding, we also noticed some functions, such as glutathione biosynthesis in Pseudomonadota or cytochrome *o* ubiquinol oxidase in Chlamydiota, to be exclusive to these specific taxa (Fig S6).

### Nitrate removal, compared with vitamin removal alone, more strongly alters the outcome of functional convergence in communities

The above findings show that complex natural communities cultured under a given, stable environment, consistently converge to similarly structured, simpler communities encoding specific metabolic functions. We hypothesised that this final functional composition mostly relates to the environmental conditions imposed during enrichment.

To test this, we designed an experiment in which replicate cultures were initiated from the *same, single* source sample and sub-cultured under three distinct medium conditions: a medium as used in the original enrichment experiments (VIT; labelled “V” in Fig. 4), the same medium but without vitamins (noVIT; “NV”), or without vitamins and nitrates (noVITnoNIT; “NVNN”) (see *Methods*) (Fig. 4A). The noVIT condition preserves the same macronutrient and electron acceptor regime while removing vitamin supplementation, whereas the noVITnoNIT condition removes nitrate as both a nitrogen source and a major electron acceptor.

**Figure 4:**
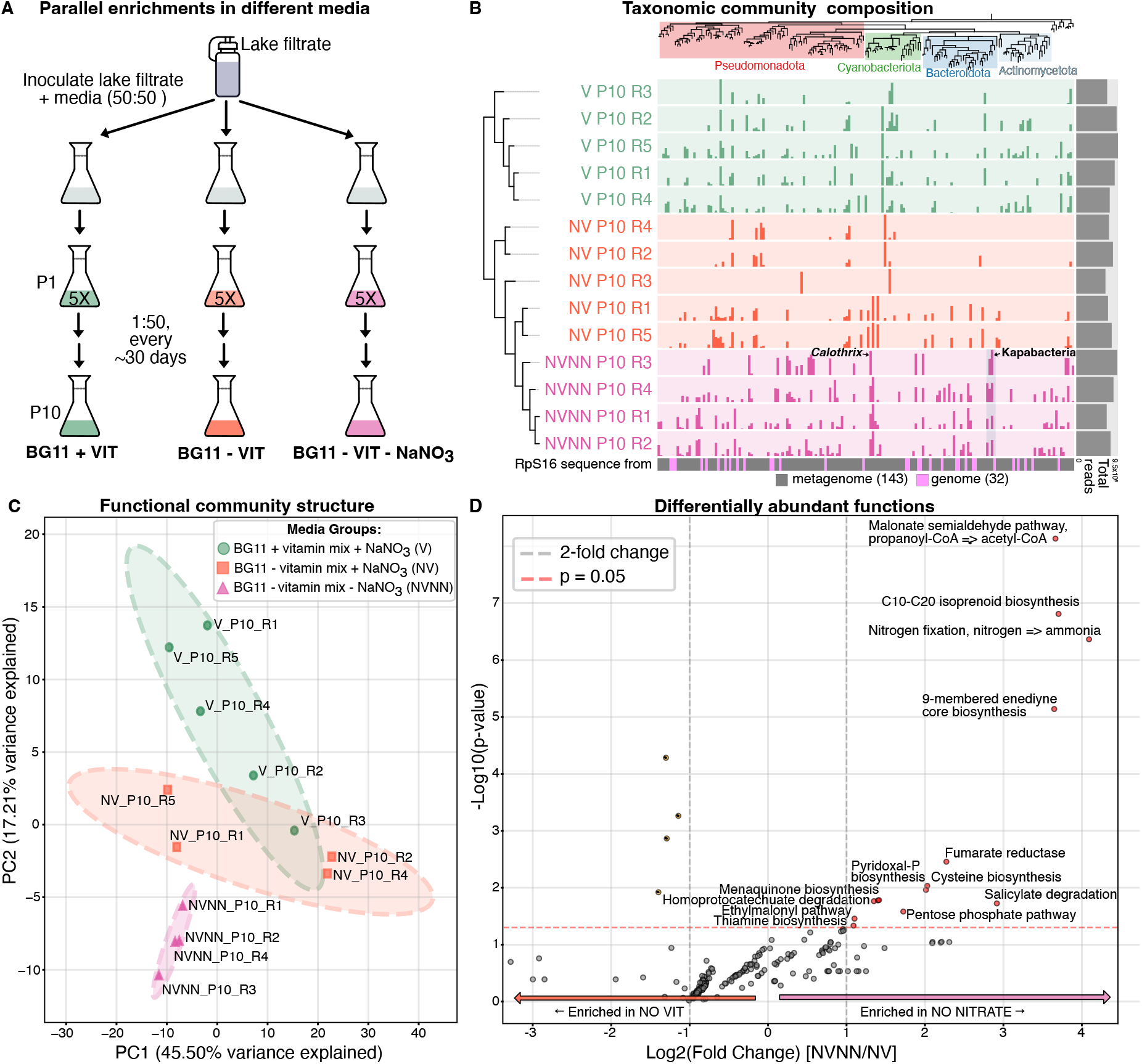
Nitrate removal rewires community composition and metabolic functional profiles. (A) Schematic workflow of the media perturbation experiment (VIT, noVIT, noVITnoNIT; five replicate lineages per condition; sub-cultured to Passage 10). Starting cultures were grown first in 50:50 lake filtrate + medium. ‘5X’ refers to replicate cultures that were grown from the starting community grown under each condition (B) Ecophylo analysis showing abundance of RpS16 sequences across the communities across the three experimental groups, indicated in green (VIT), red (noVIT), and magenta (noVITnoNIT). The dendograms to the left and top of the figure show the community sample clustering and phylogenetic relations, respectively. Some of the individual RpS16 sequences, showing clear differences among the three groups of enrichment communities are highlighted with a thick line and their name. (C) PCA based on abundance-based metric for metabolic modules, End-point communities under each group are shown as points, colored in the same way as in panel A and indicated in the legend. (D) Volcano plot showing differentially increased or decreased modules between noVITnoNIT (NVNN) and noVIT (NV). The x-axis is log_2_ fold change in the abundance-based metric across the communities in each group (NVNN/NV); dashed lines indicate 2-fold change and *p* = 0.05 thresholds. The y-axis shows *−* log_10_, FDR-corrected *p*-values.

Replicate cultures (5 for each condition) starting from the same source cultured under each media condition were sub-cultured under these media conditions for 10 sub-culture cycles (see Fig. 4A and *Methods*). Profiling of ribosomal markers showed that the taxonomic composition quickly converged to distinguishable states for each group in end-point communities, with the noVITnoNIT cultures converged to a markedly different community structure (Fig. 4B). The noVITnoNIT communities were characterised by increased relative abundance of cyanobacterial taxa including *Calothrix* (a diazotrophic cyanobacterium) and the under-studied Bacteroidata family Kapabacteria (specificaly strain PH2015), and reduced representation of multiple Pseudomonadota strains (including *Bosea, Rubrivivax*, and *Pseudomonas*) that were reproducibly abundant under nitrate-replete VIT and noVIT conditions (Fig. 4B).

As before, a similar pattern was also observed in the convergence of metabolic functional capacity. The PCA based on the abundance-based metric showed noVITnoNIT communities to form a distinct cluster separated from VIT and noVIT along PC1 (41.3% of variance; Fig. 4C). Consistent with the taxonomic shift toward diazotrophic cyanobacteria, differential module analysis between noVIT and noVITnoNIT identified *nitrogen fixation (nitrogen ⇒ ammonia)* as one of the strongest enriched functions in nitrate-free communities (Fig. 4D). In addition, nitrate-free communities showed significant enrichment of central carbon and cofactor-associated functions including the *pentose phosphate pathway, pyridoxal phosphate (vitamin B*_*6*_*) biosynthesis, menaquinone biosynthesis*, and *cysteine biosynthesis*, together with *fumarate reductase* and *salicylate degradation* (Fig. 4D). Among the most increased modules were also *C10–C20 isoprenoid biosynthesis* and *9-membered enediyne core biosynthesis* (Fig. 4D), *thiamine biosynthesis*, the *ethylmalonyl pathway*, and *homoprotocatechuate degradation* (Fig. 4D). As above, these findings indicates modules being increased in a way reflecting both imposed and emergent metabolic conditions.

## Discussion

Here, using freshwater lake samples, we analysed the impact of stable media conditions on community assembly and emergence of community function. Across eight independent enrichment experiments, initiated from temporally distinct lake communities, we observed strong and repeatable convergence of community taxonomic composition and metabolic function under carbon-free, nitrate-replete, light–dark cycling. Taxonomic convergence between initial and final communities could be detected even when using individual ribosomal genes. Despite this strong convergence across experiments, individual end-point enrichment communities showed taxonomic variance. Functional convergence - based on metabolic modules - was stronger than taxonomic convergence, with pathways for oxygenic photosynthesis, Calvin cycle carbon fixation, and nitrate-linked nitrogen metabolism consistently increasing in all enrichment communities (Table 1, Fig. 3, S6, S7). These results provide convincing experimental evidence that environmental filtering can act primarily on metabolic functions, yielding similar functional states despite species-level variability within communities. This main finding agrees with previous 16S rRNA gene amplicon-based environmental [3, 4, 9, 14] and laboratory [18, 22, 5] studies, while our metagenomics analyses provide detailed insights into the metabolic functional states.

The nitrate-replete, carbon-free, light-dark environment used here selected deterministically for photosynthesis, as well as both assimilatory nitrate reduction and dissimilatory nitrate reduction/denitrification modules, i.e. nitrate respiration (Fig. F3D). The former finding shows that phototrophs can dominate carbon-free, light-dark environments and readily sustain a community of heterotrophs around them as seen in previous studies [25, 23, 24]. The latter finding indicates nitrate serving a dual role: as a macronutrient supporting biosynthesis during growth in the light, and as an alternative electron acceptor during dark periods, when anoxic niches can emerge due to high oxygen consumption in dense cultures and under spatial structuring, which was observed in many of our unshaken cultures (Fig. 1A). Spatially structured cyanobacterial communities, including aggregates and mats are well known to result in sharp spatial and temporal oxygen gradients [24, 31, 32, 33, 34, 35]. Under such conditions heterotrophic (and some lithotrophic) lineages frequently exploit nitrate respiration, as documented in nitrate-storing sulfur bacteria and in mat communities [36, 37, 38, 33, 34, 35]. Supporting these considerations of emergent spatial structuring or enhanced respiration, resulting in increase of nitrate-related functions, we also found other metabolic functions that could be useful in such emergent niches, such as anoxygenic photosynthesis, and low-oxygen respiratory modules (e.g. *cbb*3-type cytochrome c oxidase) [39].

Consistent with the role of nitrate as a key nutrient and electron acceptor, our experiments culturing the same initial community under different media conditions showed that removing nitrate completely rewired communities, while removing vitamins had a lesser, non-significant impact. With nitrate removal communities were shifted to be dominated by diazotrophic phototrophs (e.g. *Calothrix*) and an altered metabolic profile in heterotrophs involving thiamine and menaquinone biosynthesis, functions that are associated with diazotrophic heterotroph in the rhizosphere [40, 41, 42]. Considering that no nitrogen fixation pathways are encoded in the heterotrophs enriched in nitrate-free conditions, this may suggest a symbiotic role for heterotrophs in supporting of cyanobacterial nitrogen fixation.

Vitamin limitation alone was insufficient to act as a strong driver, because key vitamin biosynthesis pathways were highly increased in all enrichment experiments here, irrespective of vitamin presence (Fig. 3). Vitamin-producing taxa and efficient vitamin recycling mechanisms were seen also in previous enrichment of cyanobacterial communities [24, 25, 43]. In natural communities, vitamin production and exchange is suggested to provide competitive advantages and buffer against periods of vitamin scarcity [44, 45, 46, 47, 48, 49, 50]. Here, however, we found deterministic enrichment of vitamin pathways even in experiments where media contained vitamins. One possible explanation could relate to bioavailability of vitamins, with *de novo* synthesis becoming important under our cultures conditions, e.g. within cyanobacterial aggregates. Another possibility is that vitamin pathways might be “hitchiking”, i.e. are co-encoded with, other increased metabolic functions such as anoxygenic photosynthesis or nitrate respiration.

A broader analysis of the taxonomic distribution of encoded metabolic functions in enriched communities suggests a reproducible, yet flexible, division of encoding across cyanobacteria and other bacteria, i.e. heterotrophs: cyanobacteria encoded oxygenic photosystems and Calvin cycle modules, whereas heterotrophs contributed to dissimilatory nitrate reduction and denitrification (Fig. F3E). Vitamin pathways (e.g., cobalamin, biotin, thiamine salvage) were consistently present but sometimes encoded in both, or either cyanobacteria or heterotrops, indicating cross-feeding in some cases and functional redundancy in others [10, 14]. Similar to photosynthesis, some functions such as thiamine salvage pathways and anoxygenic photosynthesis were consistently present in all enrichments and always encoded within heterotrophs and often monophyetically.

A striking, consistent outcome was the depletion of Actinobacteriota within enrichment communities, despite their high abundance in source communities. This pattern is consistent with field observations, where Actinobacteria are found to be enriched in free-living freshwater fractions but depleted within cyanobacterial colonies and bloom-associated aggregates [51, 52, 53]. Several factors may contribute to this observation: (i) copiotrophic conditions and photosynthetic exudates in the enrichment cultures may favor fast-growing taxa such as Proteobacteria and Bacteroidota over oligotrophic, slow-growing Actinobacteria [54, 55]; (ii) Actinobacteria may not be metabolically suited to thrive in the high nitrate medium found in enrichment conditions; or (iii) diel oscillations between high and low oxygen concentrations in denser laboratory cultures may not support the growth of strictly aerobic Actinobacterial clades. While our study was not designed to isolate these mechanisms, the reproducible decrease of Actinobacteria across our experiments may point towards general assembly rules relevant to bloom ecology [51, 52, 53, 56].

We used here latest gene-based, binning-independent approaches to achieve detailed metabolic module-level quantification across independent and replicate enrichment experiments. This allowed measurement of differential abundance of metabolic modules across enrichment vs. source communities, and their variance and their taxonomic associations within enrichment communities (Fig. S6-S7 and Fig. 3E). This revealed two groups of metabolic functions among highly differentially abundant modules, as those showing low or high variance within the enrichment communities. We hypothesise that functions with low variance - which include oxygenic photosynthesis, Calvin cycle, and assimilatory nitrate reduction - reflect “imposed niches” by a carbon-free, light-cycled, nitrate-replete environment. In contrast, the second group of selected functions with high variance - which include dissimilatory nitrate reduction/denitrification and vitamin biosynthesis - may be explained by “emergent niches” relating to microbial activities and interactions. Such emergent niches can include, for example, spatial or temporal anoxic microenvironments and vitamin or co-factor interdependencies. Modules increasing due to such emergent niches would show a recurring enrichment signature across enrichments with high replicate-to-replicate variation, as their increase may depend on community composition and local conditions within cultures.

Our results, as with prior enrichment studies [18, 22], point to tractable links between environmental inputs and community functional outputs: macronutrient/electron-acceptor regimes and physical drivers (e.g. light–dark cycling and aggregation) filter for constrained metabolic functional profiles that can be realized by alternative taxa. This suggests a trait-based engineering route for steering complex community inocula toward desired functions without bottom-up isolation [5, 57]. In our system, nitrate availability exerted a stronger and more reproducible selective effect than vitamin supplementation, consistent with a hierarchy of filters (macronutrients/electron acceptors > micronutrients) and with well-characterized redox stratification in phototrophic systems [31, 32, 33]. Phototrophic communities provide useful model systems, where future work could possibly map the boundaries of determinism by factorially varying C:N:P ratios, nitrate versus alternative electron acceptors, light intensity/spectrum, and redox fluctuation frequency, while manipulating inoculum diversity and timings of growth-dilution cycles. We expect that the metabolic pathway-level metrics described here may be useful to identify environment-function correlations over a wide parameter space even when species-level trajectories might seem variable [5, 14]. They will also help advance predicting microbial community dynamics [3, 4, 9, 5] and engineering microbial communities with desired functionality [58].

## Materials and Methods

### Sampling and culture enrichment

Lake water samples were collected from Draycote Reservoir, UK (52.317^*°*^N, 1.341^*°*^W) at multiple time points between December 2020 and October 2022. Eight enrichment cultures were established from these temporally distinct source communities: two samples collected in December 2020 from surface waters near the shore (community M 63) and from a rockpool with visible biofilm (M 67); one sample from surface waters in June 2021 (M 71); one surface water sample in May 2022 (M 77); and four rockpool samples in October 2022 (M 132, M 133, M 134, M 135). Additionally, eight source lake water samples (W 01, W 06, W 07, W 10, W 13, W 14, W 15, W 18) were collected from surface waters between April and September 2023 and filtered through 3*µ*m MCE membrane then 0.22*µ*m MCE membrane (Millipore) for direct metagenomic sequencing without culturing. Detailed methods for environmental sampling are available at [59].

Enrichment cultures were established by direct dilution of unfiltered lake water into BG11+ medium supplemented with trace vitamins (including biotin at 0.01 mg L^-1^, thiamine at 0.2 mg L^-1^, and cobalamin at 0.005 mg L^-1^) as previously described [24]. Cultures were serially passaged in 100 mL glass Erlenmeyer flasks or 150 mL medical flat bottles at low dilution factors (typically 1:10 to 1:50) under static conditions. Incubation was performed at room temperature under 12:12 hour light:dark cycles with white fluorescent illumination at 4–19 *µ*mol photons m^-2^s^-1^, measured using a LI-COR Quantum Sensor (LI-190R-BNC-5) and Light Meter (LI-250A). Enrichments were maintained for up to 12 passages with varying sub-culture periods (Fig. S1; passaged approximately every 35 days) and sampled at end-point, as well as some intermediary points for metagenomic co-assemblies (Fig. S3). Detailed methods for culturing and preservation of cyanobacterial environmental samples are available at [60].

For enrichment experiments from a single community under different nutrient environments, a single surface-water sample collected from Draycote Reservoir in May 2023 was filtered through miracloth (22– 25 *µ*m pore size) and used as the inoculum. Cultures were established under three nutritional conditions:(1) standard BG11+ with vitamin mix and nitrate (VIT; denoted V in Fig. 4), (2) BG11+ without vitamin mix (noVIT; denoted NV), and (3) BG11+ without vitamin mix and sodium nitrate (noVITnoNIT; denoted NVNN) (see *Methods* and Fig. 4A). Initial cultures, along with blank controls, were prepared by two-fold dilution of the filtered lake inoculum into 30 mL of the corresponding medium in 150 mL medical flat glass bottles. After the first growth cycle (38 days), five replicate cultures per medium were initiated alongside the original cultures. All of these cultures were then subsequently sub-cultured at 38–49 day intervals using a 1:50 dilution into fresh medium of the same composition, for 11 rounds (some cultures were lost due to biomass collapse). Prior to passaging, any aggregates or biofilms were resuspended by vigorous shaking and scraping to homogenize the inoculum. Cultures were incubated statically at room temperature under an Urban Buddy LED light panel providing white and red illumination at *∼*50 *µ*mol photons m^-2^s^-1^ with 12:12 hour light:dark cycles; temperatures periodically reached 28^*°*^C due to light panel heat emission.

### DNA extraction and metagenomic sequencing

For source lake samples, water was collected on 0.22 *µ*m MCE filters (Millipore) and stored at -80^*°*^C. Genomic DNA was extracted using the Qiagen PowerWater Pro kit following manufacturer protocols.

For enrichment cultures, endpoint communities were harvested by collecting at least 1 mL of culture suspension, centrifuging at 10,000*×g* for 5 minutes, and discarding supernatant. Wet pellet weights ranged from 36 to 124mg. Pellets were stored at -80^*°*^C prior to extraction. Genomic DNA was extracted using the Qiagen PowerSoil Pro kit with modifications as described in [24].

For enrichment experiments from a single community, but under different environments, total biomass was harvested by collecting entire culture volumes (30 mL plus 10 mL medium rinse) in pre-weighed 50 mL tubes, centrifuging at 4,000 rpm for 10 minutes, discarding supernatant, flash-freezing pellets in liquid nitrogen, and lyophilizing at -80^*°*^C using an Alpha 2-4 LD plus freeze-dryer (CHRIST, Germany). DNA was extracted from 5 mg freeze-dried biomass using the Qiagen PowerSoil Pro kit with modifications as described in [24], eluting in 50 *µ*L volume. A negative extraction control was processed alongside samples. DNA quality was assessed spectrophotometrically. Metagenomic libraries were prepared using Illumina Nextera XT and paired-end sequencing (2*×*150 bp) was performed by Novogene (Cambridge, UK) on an Illumina NovaSeq platform.

### Metagenomic assembly and binning

For both raw contigs and binned analyses, quality-filtered metagenomic reads were assembled using metaSPAdes v3.15.5 [61] using default parameters. Automatic binning was performed using the STRONG pipeline [28] for strain-resolved MAG recovery. For the analysis of all assembled contigs, MAG refinement, and high-throughput functional and metabolic annotations, we used anvi’o [62]. MAG quality was assessed using CheckM v1.2.0 [63], and only high-quality MAGs with >95% completeness and *<*5% contamination were retained for downstream analyses. Taxonomic classification of MAGs was performed using GTDB-Tk v2.3.0 [64] against the Genome Taxonomy Database (GTDB) release 214 [65].

### High-resolution community profiling using ribosomal proteins

To gain comprehensive insights into taxonomic diversity captured by the metagenomic assemblies beyond the recovered genomes, we used the anvi’o EcoPhylo workflow [29] with both ribosomal protein L19 (RpL19) and Ribosomal protein S16 (RpS16). Briefly, the EcoPhylo workflow (1) identified all RpS16 and/or RpL19 sequences in assembled contigs using the Pfam model PF01245 (for RpL19) and PF00380 (for RpS16) with HMMER v3.3.2 [66], (2) clustered them at 99% nucleotide identity using MMseqs2 v13.45111[67], (to generate a non-redundant set of ribosomal protein sequences that represent distinct taxa, (3) performed a read recruitment analysis using the representative sequences from each cluster to estimate the detection and coverage of each taxon across metagenomes using Bowtie2 v2.4.5 [68], and (4) calculated a phylogenetic tree using FastTree [69] to visualize evolutionary relationships between taxa based on the final ribosomal sequences that represent them. We then used the anvi’o program `anvi-run-scg-taxonomy` to estimate the taxonomic affiliation of the final EcoPhylo representative sequences using the GTDB prior to visualization of the RpS16 / RpL19 diversity and biogeography.

### Functional annotation and metabolic reconstruction

To quantify community metabolic potential independently of MAG recovery, we performed module-level functional analyses directly on all assembled contigs in a given culture. For this, we used the anvi’o program anvi-run-kegg-kofams [70] to identify KEGG Orthologs (KO) in assembled contigs against the KEGG database [71] (release 2023-10-15), and used the program `anvi-estimate-metabolism` [30] to estimate presence and completion of metabolic modules in our data. For downstream analyses, we only retained metabolic modules with ‘pathwise’ completeness score of ≥0.75, following KEGG recommendations [72].

We extracted two complementary metrics to assess functional enrichment of metabolic modules across communities. First, a coverage-based metric was obtained using the “pathwise module completeness” output from anvi-estimate-metabolism [30]. This metric calculates the fraction of essential enzymes (KOs) annotated in a sample, taking the maximum across all possible enzymatic paths through the module definition. For each module, the average module coverage was computed by summing the read coverages of all contigs carrying KOs belonging to that module, divided by the total number of KOs present in the module. This reflects the average sequencing depth invested in each metabolic pathway per metagenome. Modules with pathwise completeness ≥75% were considered present in a sample.

Second, we calculated a count-based metric to assess module abundance normalized by community richness. For this, we used the “stepwise copy number” output from anvi-estimate-metabolism, which represents the minimum copy number across all top-level steps in a module definition (see [30] for technical details). To correct for differences in microbial diversity between samples, we normalized stepwise module copy numbers by dividing them by the number of distinct ribosomal proteins sequence clusters detected in each metagenome. This metric represents the expected module copy number per coexisting microbial population.

For differential analyses, sample*×*module matrices were constructed for both metrics. Modules absent from all samples in both source and enrichment groups were excluded. For each module, log_2_ fold changes were calculated as the ratio of the mean abundances in the enrichment to source communities, and statistical significance was assessed using two-sided Student’s *t*-tests on log_10_(*x*+0.001)-transformed values. Raw *p*-values were corrected for multiple testing using the Benjamini–Hochberg false discovery rate (FDR) procedure. Modules were considered significantly enriched if FDR < 0.05 and absolute log_2_ fold change > 0.5.

To examine metabolic encoding across cyanobacteria and heterotrophs, we mapped each module to contributing MAGs based on pathwise completeness ≥0.75. For each module in each enrichment sample, we summed, separately the relative abundances of cyanobacterial versus heterotrophic MAGs that encode module and then expressed this as percentage of total abundance of all MAGs encoding that module. MAGs were classified as cyanobacterial or heterotrophic based on GTDB phylum-level assignment (Cyanobacteriota vs. all other phyla). Statistical significance of deviation from equal contribution (50:50) was assessed using one-sample *t*-tests across the eight enrichment replicates.

### Statistical analysis of community convergence and functional enrichment

All statistical analyses were performed using custom Python scripts leveraging scikit-learn v1.2.2 [73], pandas v2.0.1 [74], and NumPy v1.24.3 [75].

Principal component analysis (PCA) was performed on centered and scaled taxonomic (RpL19-based relative abundances) and functional (KEGG module relative abundances) matrices using scikit-learn’s PCA implementation. Module loading contributions to principal components were calculated as the correlation between module abundances and PC scores. Variance explained by each principal component was calculated from eigenvalues of the covariance matrix. Statistical significance of group separation between source and enrichment communities, as well as treatment groups in media manipulation experiments, was assessed using PERMANOVA implemented using the skbio.stats.distance.permanova function, employing permutation tests (n=999) on Euclidean distances between group centroids.

Coefficient of variation (CV) for each module across enrichment replicates was calculated as the ratio of standard deviation to mean relative abundance (*CV* = *σ/µ*). Modules were classified as having low (CV < 0.35), intermediate (0.35 *≤* CV < 1.0), or high (CV ≥ 1.0) variance, based on the distribution of all CV values.

All visualizations were generated using Matplotlib v3.7.1, Seaborn v0.12.2, or the anvio interactive interface. Statistical significance thresholds were set at *α* = 0.05 for all tests unless otherwise specified.

## Author Contributions

SJND, AS, and OSS conceptualised and developed the methodology of the study. SJND, MC, and AS performed the sampling and experiments. AS, ME, and OSS performed formal analysis and visualisation of the data. SJND, AS, ME, and OSS wrote the manuscript and completed review and editing. OSS obtained funding, and OSS and MC were responsible for project administration.

## Aknowledgements

We acknowledge the members of Soyer Lab (https://warwick.ac.uk/fac/sci/lifesci/research/osslab/) and Meren Lab (https://merenlab.org) for discussions about the analyses of data and presentation of results. We acknowledge cooperation and support of Keiron Maher and Gemma Hill at Severn Trent with sampling permissions at Draycote Water, and Mark Dunkley, Andy Felthouse, Matt Rowley, and David Rowe at the Draycote Water Sailing Club and Jerko Rosko (former member of Soyer Lab) with sample collection.

