## Supplementary Information for "Enrichment of convergent metabolic functions in microbial communities through imposed and emergent environmental niches"

February 11, 2026

### List of Tables

S1 Assembly and annotation statistics for metagenomic samples . . . . . . . . . . . . . . . . . . . 3

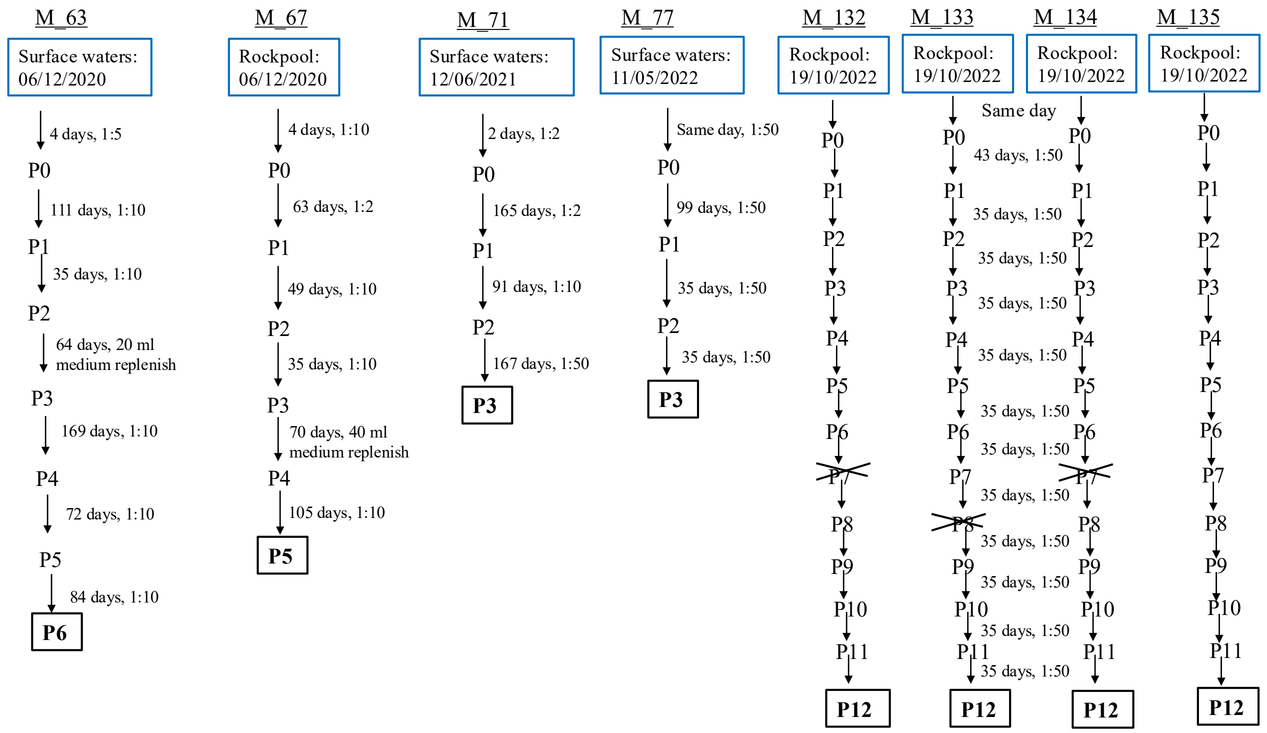

**Figure S1: Sub-culturing regimes for enrichment communities established from temporal lake samples.** Lake water samples (blue boxed details) were collected either from the lake water near the shore or from rockpools at diverse locations (see Methods). 'P' represents passage number, with P0 as the original cultures that were inoculated in BG11+ vitamin mix medium with dilution from original samples, either on the same day or after a short period of lab incubation. Sub-culture passages and culture durations are indicated on the figure, together with the numbers of days and the dilution factor into fresh medium used. In a few cases, medium was replenished due to culture evaporation. The boxed final passage numbers represent the final characterised communities, with sample IDs shown in the headings. Cultures were grown in 100 ml Erlenmeyer culture flasks or in 150 ml medical flat glass bottles, typically with 30 ml final volumes. From the October 2022 samples, passage numbers crossed out represent community collapse. In these cases, sub-culturing was repeated from the previous passage sample, again with a 1:50 dilution.

**Table S1:** Assembly and annotation statistics for metagenomic samples

| Sample | Total Reads | Total Length (bp) | Num Contigs | N50 (bp) | L50 | Longest Contig (bp) | Num Genes |
|---|---|---|---|---|---|---|---|
| M_132 | 93,027,578 | 280,582,668 | 378,593 | 1,140 | 34,245 | 957,661 | 534,866 |
| M_133 | 82,072,452 | 331,701,172 | 446,491 | 1,211 | 41,411 | 1,292,358 | 631,329 |
| M_134 | 77,798,192 | 424,112,571 | 493,369 | 1,751 | 39,170 | 1,258,215 | 706,382 |
| M_135 | 84,805,046 | 375,613,587 | 340,520 | 4,276 | 9,085 | 1,778,518 | 584,947 |
| M_63 | 79,494,758 | 180,146,349 | 233,813 | 1,454 | 15,544 | 853,854 | 344,584 |
| M_67 | 109,529,224 | 494,027,024 | 667,849 | 1,442 | 47,187 | 844,049 | 957,233 |
| M_71 | 101,187,226 | 411,479,467 | 462,146 | 1,831 | 30,907 | 2,625,472 | 741,987 |
| M_77 | 97,812,882 | 270,729,295 | 372,810 | 1,150 | 27,874 | 1,265,934 | 520,613 |
| W_01 | 74,402,146 | 1,120,498,306 | 2,539,959 | 495 | 451,436 | 343,579 | 3,063,818 |
| W_06 | 79,619,144 | 1,226,804,826 | 2,856,204 | 479 | 485,937 | 866,479 | 3,411,117 |
| W_07 | 95,653,372 | 1,175,852,193 | 2,727,618 | 524 | 412,495 | 459,514 | 3,246,909 |
| W_10 | 67,782,306 | 1,005,514,782 | 2,263,475 | 513 | 368,127 | 878,920 | 2,711,121 |
| W_13 | 77,593,422 | 1,306,492,769 | 2,981,320 | 492 | 528,007 | 275,199 | 3,578,314 |
| W_14 | 74,498,880 | 1,274,169,250 | 2,893,896 | 497 | 522,344 | 317,484 | 3,371,101 |
| W_15 | 140,306,802 | 1,620,992,101 | 3,330,017 | 662 | 436,206 | 586,040 | 4,123,266 |
| W_18 | 70,840,458 | 1,373,206,263 | 3,247,107 | 454 | 624,463 | 276,061 | 3,690,075 |
| NVNN_P10_R1 | 69,172,890 | 257,333,830 | 278,210 | 2,164 | 10,823 | 2,130,088 | 452,161 |
| NVNN_P10_R2 | 78,128,452 | 271,473,104 | 269,887 | 4,258 | 4,954 | 1,146,810 | 454,584 |
| NVNN_P10_R3 | 93,279,874 | 132,498,956 | 49,776 | 59,047 | 442 | 1,557,922 | 156,561 |
| NVNN_P10_R4 | 84,707,242 | 264,728,551 | 152,808 | 38,454 | 1,155 | 1,569,995 | 353,481 |
| NV_P10_R1 | 72,231,184 | 326,519,114 | 494,331 | 1,053 | 42,835 | 2,786,513 | 622,757 |
| NV_P10_R2 | 83,539,720 | 168,659,284 | 165,948 | 13,216 | 2,496 | 232,005 | 294,420 |
| NV_P10_R3 | 66,423,224 | 166,583,483 | 44,150 | 22,390 | 1,800 | 1,480,983 | 240,860 |
| NV_P10_R4 | 74,596,548 | 206,405,250 | 247,572 | 3,780 | 6,352 | 903,009 | 376,428 |
| NV_P10_R5 | 80,619,888 | 229,244,327 | 248,568 | 1,865 | 13,724 | 1,100,890 | 378,243 |
| V_P10_R1 | 87,478,708 | 355,354,589 | 298,043 | 5,161 | 9,411 | 1,343,699 | 555,388 |
| V_P10_R2 | 91,879,908 | 445,168,834 | 596,849 | 1,667 | 35,330 | 1,156,722 | 833,353 |
| V_P10_R3 | 70,358,094 | 177,256,850 | 207,803 | 10,427 | 2,776 | 799,391 | 329,652 |
| V_P10_R4 | 76,178,726 | 380,304,987 | 477,695 | 1,657 | 25,121 | 1,316,469 | 711,132 |
| V_P10_R5 | 94,889,976 | 373,178,139 | 530,951 | 1,114 | 32,237 | 1,156,722 | 691,855 |

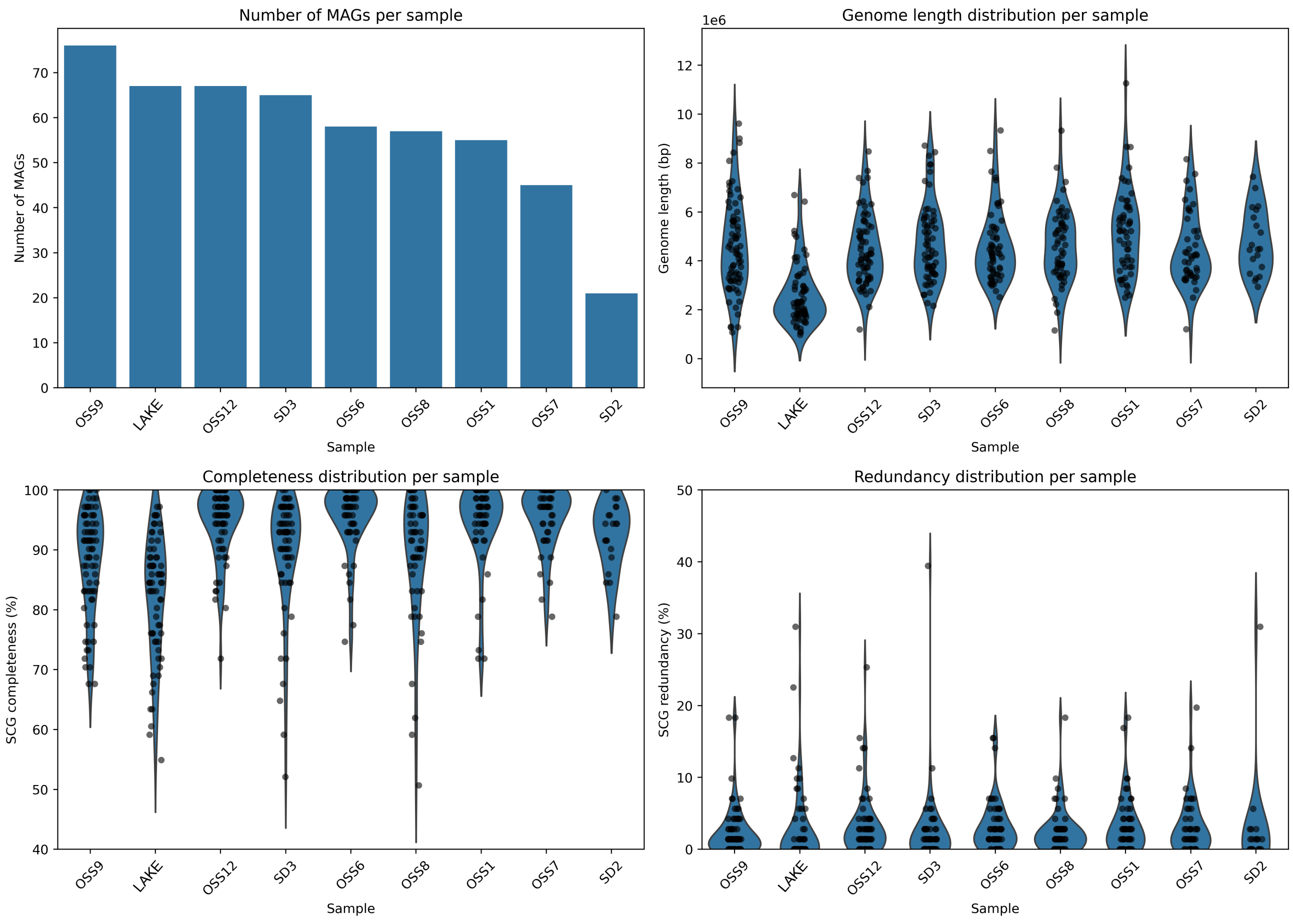

**Figure S2: Metagenome-assembled genome (MAG) recovery and quality metrics from STRONG co-assemblies**. Each co-assembly combines multiple individual enrichment culture samples at different passage numbers: OSS1 (M_74, M_77), OSS12 (M_68, M_71), OSS6 (M_81, M_85, M_134), OSS7 (M_80, M_84, M_133), OSS8 (M_79, M_83, M_132), OSS9 (M_82, M_86, M_135), SD2 (M_62, M_63), SD3 (M_64, M_67), LAKE (W_06, W_10, W_13, W_15). Panels show: (A) total number of MAGs recovered per co-assembly, (B) genome size distributions, (C) completeness estimates, and (D) redundancy estimates as determined by single-copy core genes (SCGs) obtained from *anvi-estimate-genome-completeness*

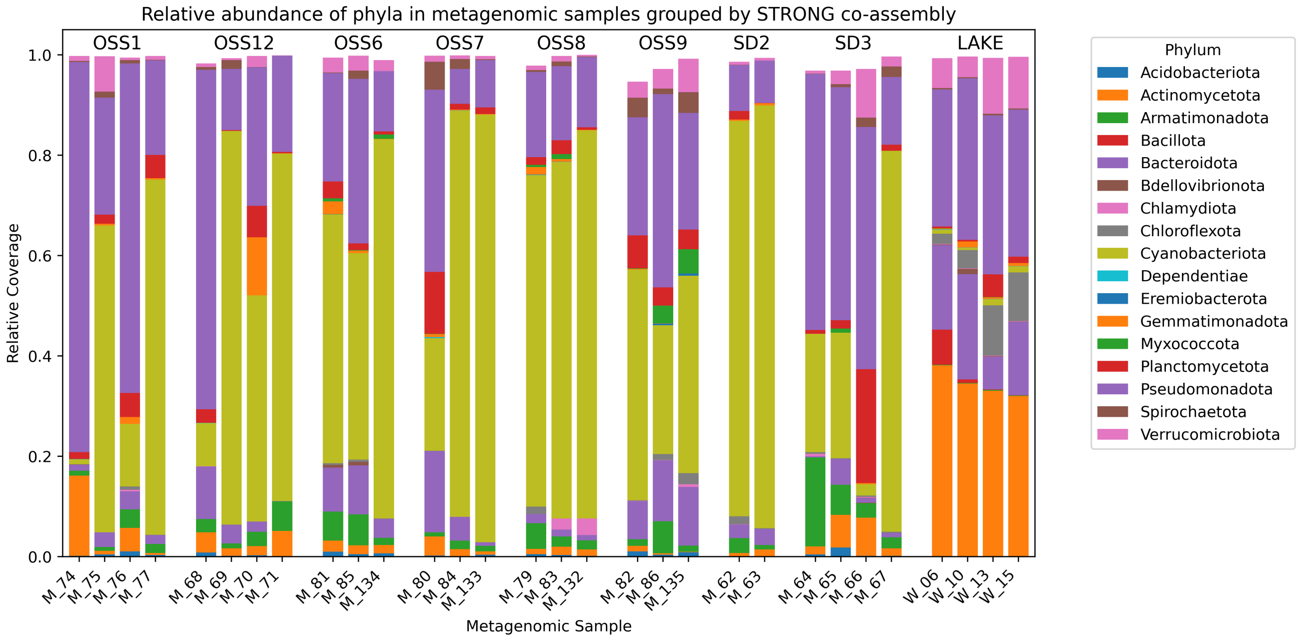

**Figure S3: Relative coverage of high-quality metagenome-assembled genome obtained from STRONG co-assemblies.** Each co-assembly combines multiple individual enrichment culture samples at different passage numbers: OSS1 (M_74, M_77), OSS12 (M_68, M_71), OSS6 (M_81, M_85, M_134), OSS7 (M_80, M_84, M_133), OSS8 (M_79, M_83, M_132), OSS9 (M_82, M_86, M_135), SD2 (M_62, M_63), SD3 (M_64, M_67), LAKE (W_06, W_10, W_13, W_15). Taxonomy inferred from the Bacteria_71 single copy gene (SCGs) using *anvi-estimate-scg-taxonomy*.

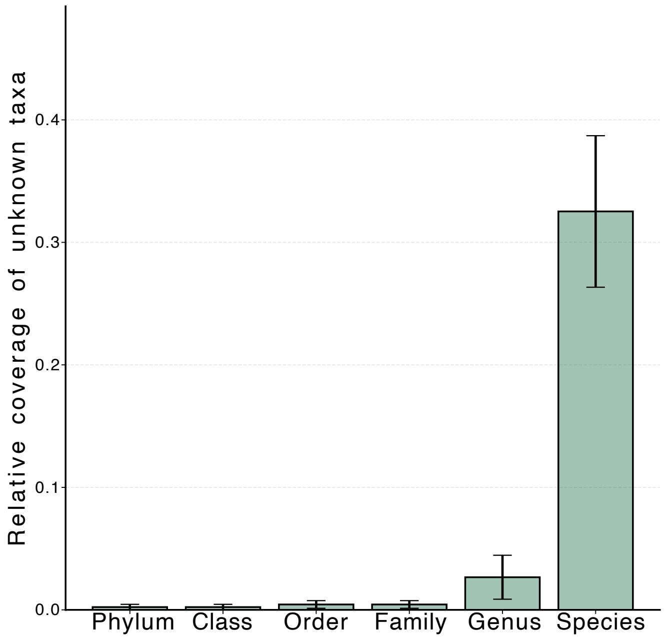

**Figure S4: Relative coverage of taxonomically unassigned RpL19 lineages across taxonomic ranks.** RpL19 marker sequences (whose coverage was greater than 0) from all ENRICHMENT samples were obtained from the ecophylo analysis presented in Figure 2 and classified using anvi-run-scg-taxonomy against the GTDB reference database (v2.3.0). Bars show, for each rank (phylum to species), the fraction of total RpL19 coverage that remained unassigned (“unknown”), providing a rank-resolved measure of taxonomic assignment uncertainty.

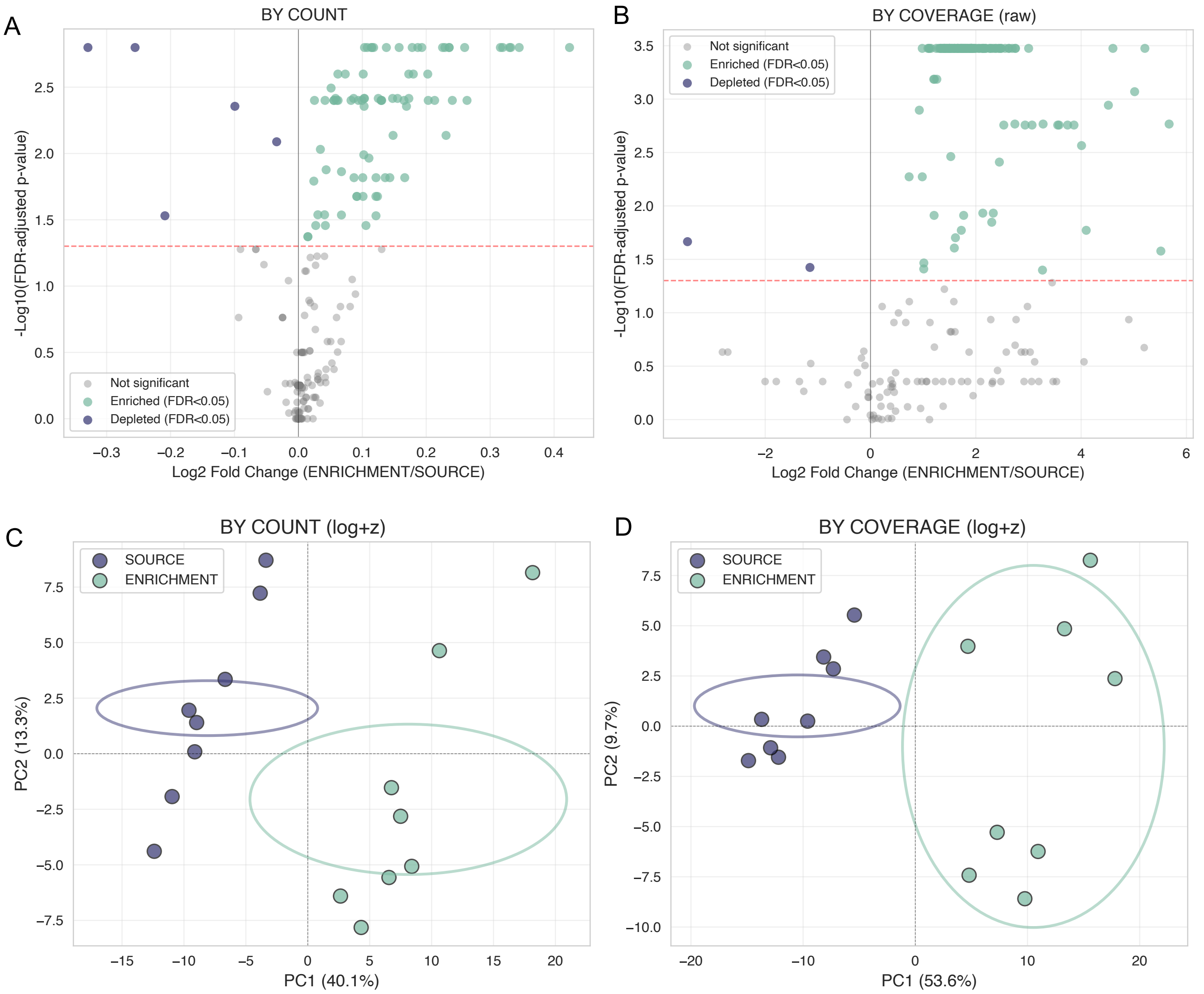

**Figure S5: Functional metabolic profile comparison between source and enrichment samples**. Differential abundance analysis of metabolic genes between source environmental samples and enrichment cultures using (A) module count-based and (B) coverage-based metrics. Volcano plots show log2 fold change (ENRICHMENT/SOURCE) versus statistical significance (FDR-adjusted p-value ¡ 0.05). Green points indicate KEGG modules significantly enriched in cultures, blue points indicate genes depleted in cultures. Principal component analysis (PCA) of metabolic profiles using log-transformed and z-scored data for (C) module count-based and (D) coverage-based metrics (see *Methods* for details), showing clear separation between source (blue) and enrichment (green) communities. Ellipses represent 95% confidence intervals.

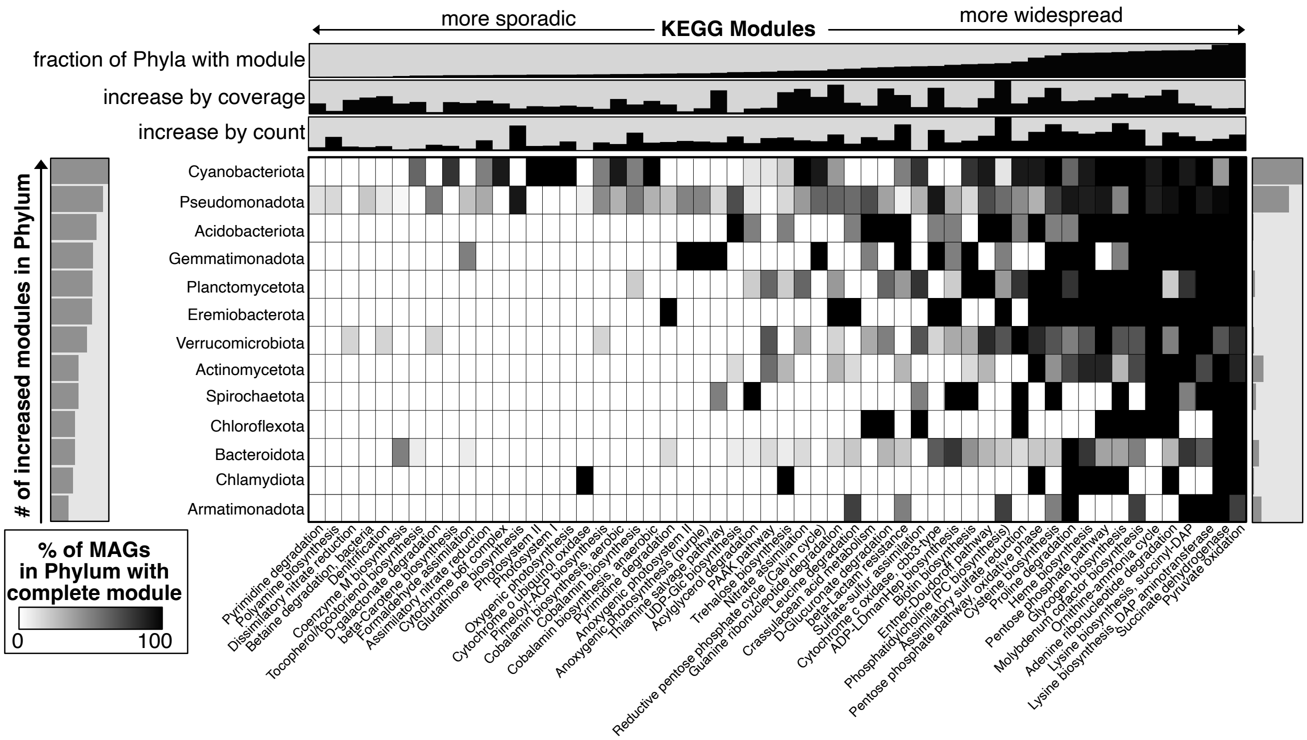

**Figure S6: Distribution of modules, which are significantly increased in the enrichment communities, across metagenome-assembled genomes (MAGs) found in those communities, collapsed at the phylum level.** MAGs included 87 high-quality MAGs recovered from ENRICHMENT samples collapsed at the phylum level. Each row represents a phylum, each column a KEGG module, and filled cells indicate the fraction of MAGs in that phylum with module completeness $\geq 0.75$. Rows are ordered according to the phyla containing the largest number of enriched modules and columns are ordered by the fraction of Phyla containing each module.

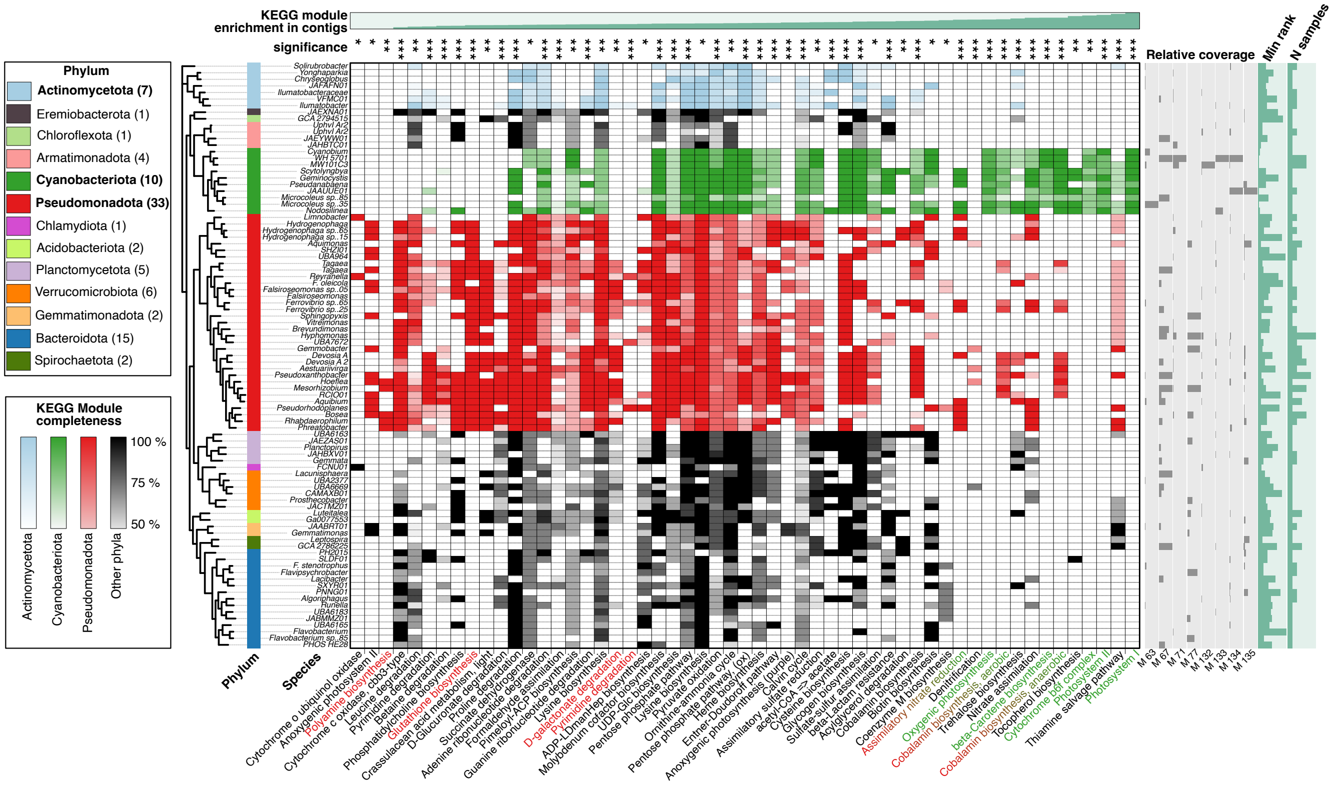

**Figure S7: Distribution of modules, which are significantly increased in the enrichment communities, across metagenome-assembled genomes (MAGs) found in those communities.** The heatmap displays KEGG module completeness (color intensity: 50-100% complete) for 87 high-quality MAGs recovered from ENRICHMENT samples and arranged by phylogenetic classification (Rpl19 sequences). MAGs are grouped by phylum with counts shown in parentheses: Pseudomonadota (33), Bacteroidota (15), Cyanobacteriota (10), Actinomycetota (7), Verrucomicrobiota (6), Planctomycetota (5), Armatimonadota (4), Gemmatimonadota (2), Acidobacteriota (2), Spirochaetota (2), and singleton phyla Chlamydiota, Chloroflexota, and Eremiobacterota. The three phyla that explain the greatest variation between source and enrichment culture composition — Cyanobacteriota, Pseudomonadota, and Actinomycetota — are color-coded in green, red, and light blue, respectively. Relative coverage bar charts (right) show MAG abundance distributions across individual enrichment samples (M_63, M_67, M_71, M_77, M_132, M_133, M_134, M_135). Min rank and N samples bar charts indicate the minimum rank of the MAG in each sample (based on coverage of MAGs) and the number of samples in which each MAG was detected, respectively. The top panel displays metabolic module increase (by coverage) in cultures relative to source samples (log2 fold change color scale), with asterisks denoting statistical significance (FDR ¡ 0.05). Metabolic modules are ordered by increasing module increase from left to right.

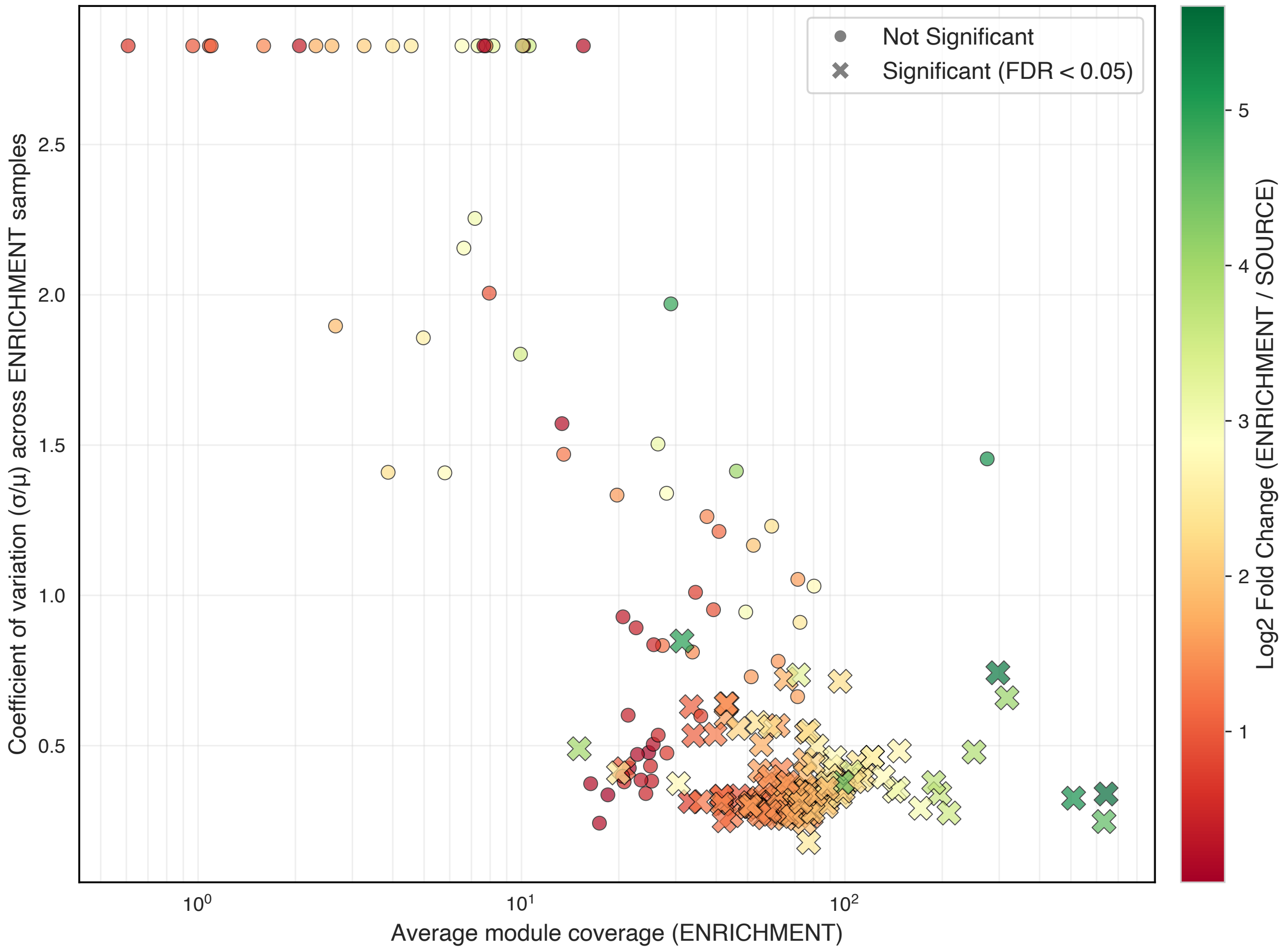

**Figure S8: Relationship between metabolic module abundance, variability, and enrichment in cultures**. Scatter plot showing the metabolic module abundance in enrichment communities (x-axis) and abundance coefficient of variation (CV = $\sigma/\mu$) across those communities (y-axis). Each point indicates a module. Points are colored by the log2 fold change of a module between enrichment and source communities (ENRICHMENT/SOURCE), with warmer colors (red-orange) indicating depletion in cultures and cooler colors (yellow-green) indicating enrichment. Circles represent modules with non-significant differential abundance, while X markers indicate modules significantly different between source and enrichment samples (FDR $<$ 0.05). High-abundance modules tend to show lower variance across replicates, while low-abundance modules display greater variability.

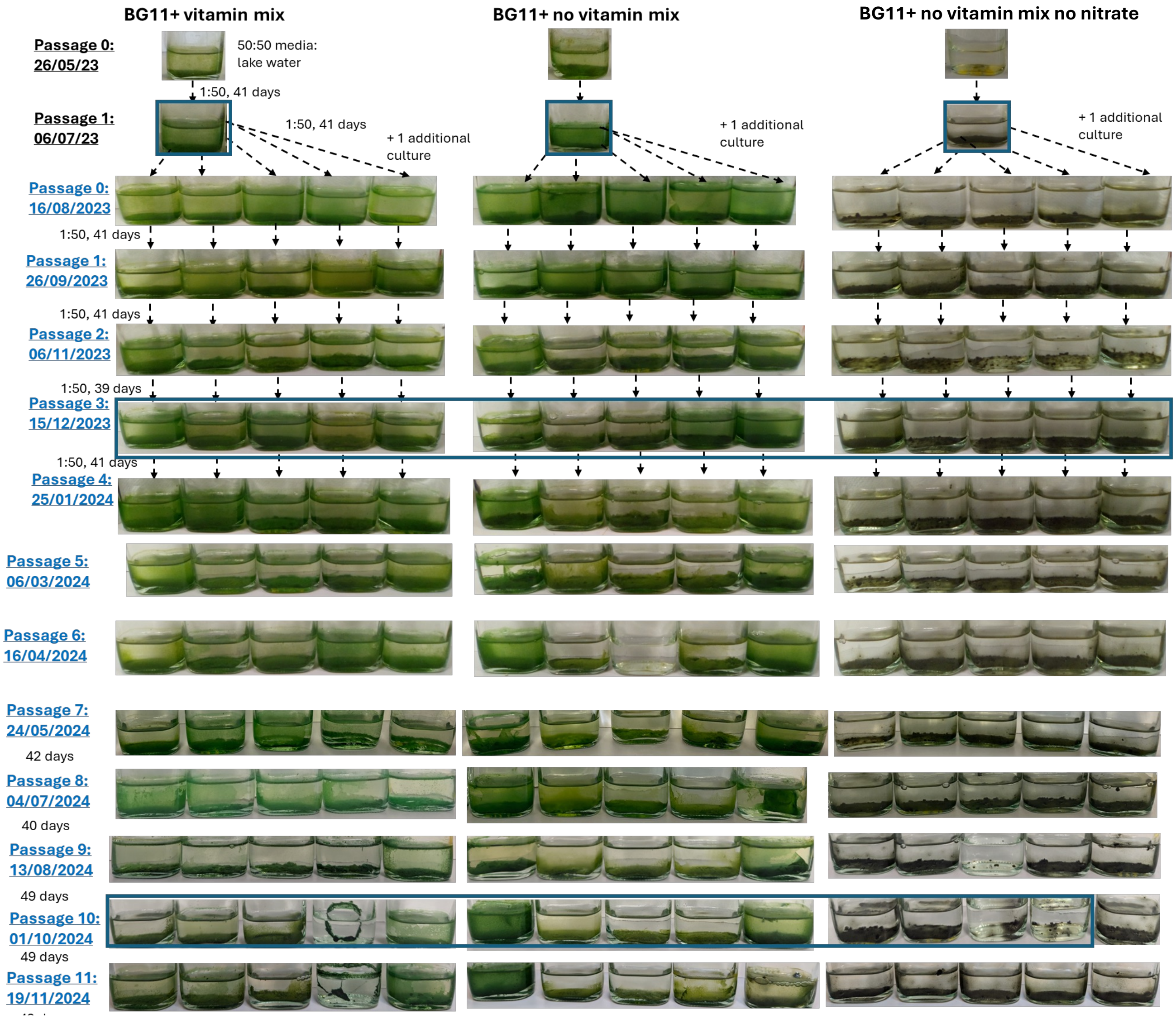

**Figure S9: Enrichment culture development under different medium compositions.** Three parallel enrichment series were established from the same initial inoculum (Passage 0: 26/05/23, 50:50 lake water:media) and maintained under varying nutritional conditions: BG11+ with vitamin mix (left), BG11+ without vitamin mix (center), and BG11+ without vitamin mix or nitrate (right). Cultures were serially passaged at 1:50 dilution with transfer intervals of 39-49 days. Blue boxes indicate passages selected for shotgun metagenomic sequencing (Passage 3: 15/12/2023 and Passage 10: 01/10/2024).
